# Failure of vergence size constancy challenges our understanding of visual scale

**DOI:** 10.1101/2020.07.30.228940

**Authors:** Paul Linton

## Abstract

The closer an object is, the more the eyes have to rotate to fixate on it. This degree of eye rotation (or vergence) is thought to play an essential role in size constancy, the process of perceiving an object as having a constant physical size despite changes in distance. But vergence size constancy has never been tested divorced from confounding cues such as changes in the retinal image. We control for these confounding cues and find no evidence of vergence size constancy. This has three important implications. First, we need a new explanation for binocular vision’s contribution to visual scale. Second, the vergence modulation of neurons in V1 can no longer be responsible for size constancy. Third, given the role attributed to vergence in multisensory integration, multisensory integration appears to be more reliant on cognitive factors than previous thought.

## Introduction

The Taylor illusion (1) provides an important insight into how we see objects as having a constant physical size despite changes in distance altering the size of the retinal image: An observer views an afterimage of their hand in complete darkness. When the observer moves their hand towards themselves the hand appears to shrink, whilst when the observer moves their hand away, the hand appears to grow. Because afterimages fix the retinal image, any change in the perceived size of the hand must be due to extra-retinal influences. One initial interpretation is that this effect is due to multisensory integration, with the proprioceptive signal from the physical hand influencing the perceived size of the visual hand. However, recent work has shown that the Taylor illusion is almost entirely due to the changing gaze position of the observer as their eyes track the motion of their hand in darkness (2)(3). This coheres with over a century’s worth of experimentation confirming vergence (the angular rotation of the eyes) as one of our most important cues for size constancy (4)(5), and the neurophysiological finding that vergence modulates the vast majority of neurons in V1 (6).

However, one surprising fact is that to the best of our knowledge vergence has never been tested as a cue to perceived size divorced from confounding cues (such as changes in the retinal image or changes in hand position) which inform the observer about the change in distance. The reason for this is easy to appreciate. In order to drive vergence, we either have to present diplopia in the retinal image, typically from the retinal slip of an LED moving in depth, or ask the observer to track their own hand. However, hand movements directly inform the observer about changes in distance, whilst above threshold diplopia can also provide absolute distance information (7).

We tested vergence size constancy using parameters that were consistent with the Taylor illusion in (2)(a 3° visual target moving in depth from 50 to 25cm over 5 seconds), but in piloting were found to minimise subjectively noticeable retinal slip, and provided no proprioceptive distance information. On each 5 second trial we increased or decreased the physical size of a 3° visual target over the 5 seconds (by up to +/−20%), whilst at the same time increasing the vergence distance of the target from 50 to 25cm. The question was whether the change in vergence would have any effect on participants’ judgements of whether the target got bigger or smaller?

## Results

We found no evidence that vergence affects size judgements. A hierarchical Bayesian psychometric function is fitted using the Palamedes ToolBox (8) (with the toolbox’s standard priors) (Fig.1A) in order to provide a population level estimate of the bias from vergence on size judgements (Fig.1B). What we find at the population level is a negligible non-significant bias in the wrong direction for size constancy (–0.219%; 95% CI: –1.82% to 1.39%). From a Bayesian perspective, we performed a JZS Bayes factor (9), and the estimated Bayes factor (3.99, ±0.03%) suggests that the data are four times more likely under the null hypothesis (bias = 0) than under the alternative hypothesis (bias ≠ 0). From a frequentist perspective, we performed an inferiority test (10), taking the detection threshold for our most sensitive observer (1.43%) as the smallest effect size of interest. Since we have a directional hypothesis, we perform an inferiority test by taking the 90% confidence interval of the population bias in the predicted direction (0.96%). Since this is smaller than 1.43%, and therefore undetectable by even our most sensitive observer, from a frequentist perspective we can conclude that any vergence size constancy effect is effectively equivalent to zero.

**Figure 1.**
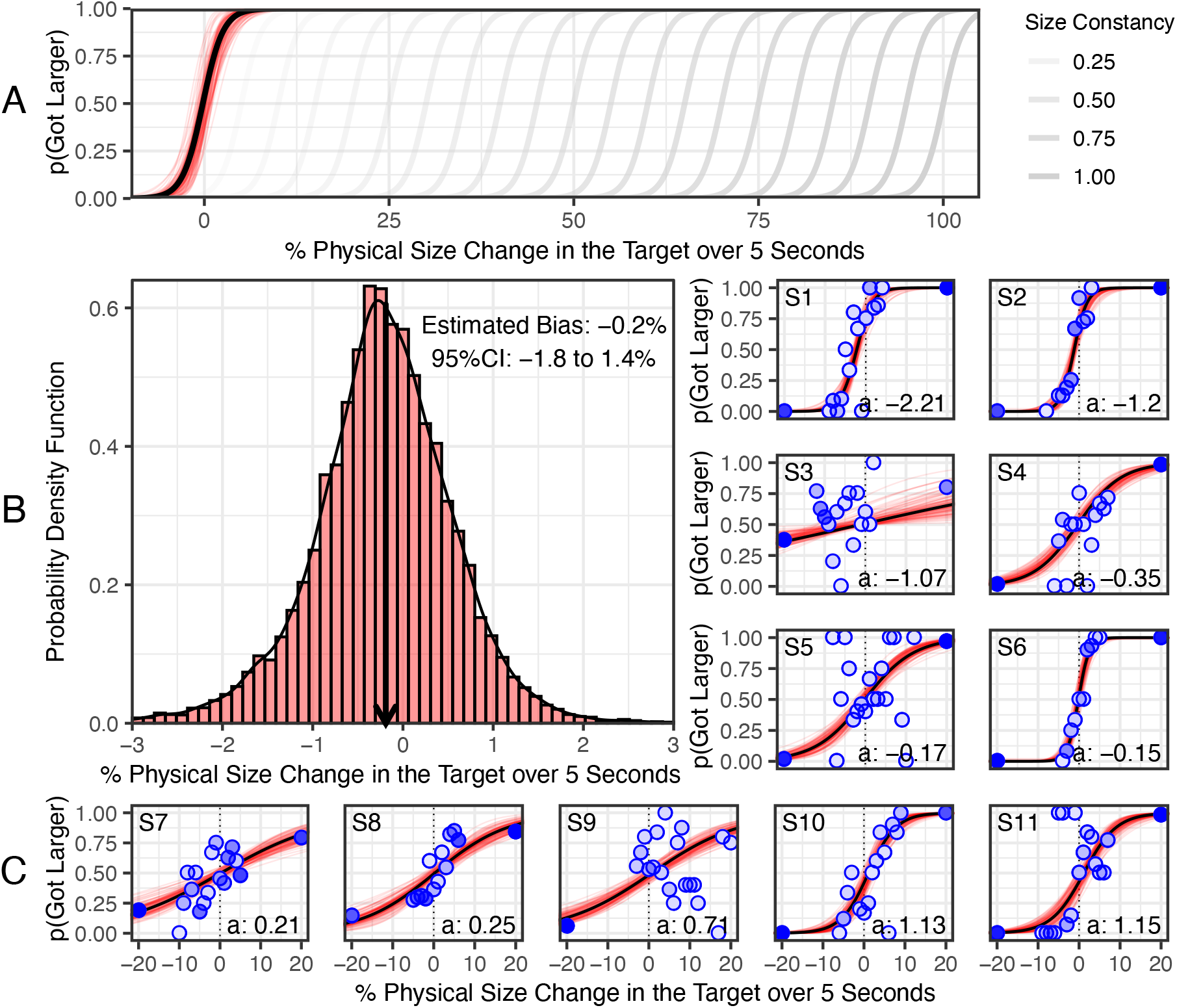
Experimental results of larger/smaller judgements against physical size change indicate no evidence of vergence size constancy. A. Hierarchical Bayesian population level psychometric function in black (based on 15,000 posterior estimates, 100 representative estimates in red), plotted against predictions for various vergence size constancy effect sizes (in grey). B. Histogram of the 15,000 posterior estimates of the population bias indicating a negligible non-statistically significant bias in the wrong direction for vergence size constancy. C. Individual results, with blue dots indicating physical size changes tested (darkness indicates frequency), and Bayesian psychometric function fitted in black (based on 15,000 posterior estimates, 100 representative estimates in red).

## Discussion

These results have three important implications. First, these results challenge our understanding of how binocular vision affects visual scale. Helmholtz illustrated the importance of binocular vision for size perception by demonstrated that the altering the interpupillary distance drastically affects the perceived size of a real-world scene (11)(12). This effect is typically attributed to two aspects of binocular vision: vergence and vertical disparities (12). However, our results challenge vergence as a cue to visual scale, whilst the experimental literature has found vertical disparities to be effective in only very limited contexts (13). By a process of elimination this leads to the conclusion that horizontal disparities may instead be responsible for binocular vision’s contributions to visual scale (horizontal disparities fall off with distance^2^, and so a scene may look miniature if the distribution of horizontal disparities is consistent with the scene being closer).

Second, V1 reflects the perceived rather than retinal size of objects (14). The finding that the vast majority of neurons in V1 are responsive to changes in vergence appeared to provide a neurophysiological underpinning for binocular size constancy in V1 (6). Recent work supports this hypothesis (15), although posits the need for recurring processing within V1. By contrast, our results suggest that the vergence modulation recorded in V1 cannot be responsible for size constancy, and instead suggests that size constancy is much more reliant (and indeed may be wholly reliant) on top-down processing than previously thought.

Third, we need a new explanation for the Taylor illusion. The changing size of an afterimage of the hand is almost entirely governed by vergence when vergence and the hand are moved in opposite directions (2), and yet we find no effect of vergence on percieved size when the vergence signal is isolated. The key difference is that those previous studies provided participants with subjective knowledge about their changing gaze position, either through proprioceptive signals (hand movement) or through the consciously percieved motion in depth of an LED (when hand and gaze are moved in opposite directions). This suggests that the Taylor illusion is largely a cognitive phenomenon, reliant upon participants having subjective knowledge about their gaze position, rather than reflecting a fundamental mechanism of multisensory integration.

## Methods

Participants viewed two 3° targets (H rotated 90°) separated laterally in a stereoscope with parallax barriers (so the left eye saw the right target and the right eye saw the left target), and vergence was increased or decreased by increasing or decreasing the separation between the targets. Vergence-accommodation conflict was kept within reasonable bounds (+/− 1 dioptre) by fixing accommodation at 33cm using contact lenses. Stimuli were presented on an OLED display (ensuring no residual luminance from black pixels) using PsychToolBox (16), and distortions of the target with lateral movements on the display were controlled using OpenGL. Participants viewed the stimuli in complete darkness through red filters, with blackout fabric ensuring that any residual luminance was eradicated.

Participants completed 200 trials using maximum likelihood estimation to specify the size change on each trial ((17)(18)). For the purposes of an inferiority test (10), we wanted to demonstrate that any size change was so small that it couldn’t be detected by our most sensitive observer. During piloting the smallest detection threshold was found to be a 1.5% size change, and given plausible assumptions about detection thresholds (5%) and lapse rate (2%), estimated we would achieve this degree of precision with 5+ observers. 11 observers participated (8 female, 3 male; age ranges 20-34, average age 24.5), the author and 10 participants recruited online (3 more participants were initially recruited but 1 was excluded because they could not fuse the target, and 2 were excluded because they could not get clear vision with the contact lenses). All participants were screened to ensure accommodation, vergence, and stereoacuity were normal, and the study was approved by the School of Health Sciences Research Ethics Committee at City, University of London. Experimental code, data, and analysis, are openly available at: https://osf.io/5nwaz/

## Acknowledgements

Thank you to Christopher Tyler, Simon Grant, Joshua Solomon, Matteo Lisi, Byki Huntjens, Michael Morgan, Chris Hull, and Pete Jones for advice, and Salma Ahmad, Mark Mayhew, Deanna Taylor, Jugjeet Bansal, and the CitySight clinic for assistance with contact lens.

